# Rapid quantitative imaging of high intensity ultrasonic pressure fields

**DOI:** 10.1101/2020.02.15.951046

**Authors:** Huiwen Luo, Jiro Kusunose, Gianmarco Pinton, Charles F Caskey, William A Grissom

## Abstract

High-intensity focused ultrasound (FUS) is a noninvasive technique for thermal or mechanical treatment of tissues that can lie deep within the body, with a growing body of FDA-approved indications. There is a pressing need for methods to rapidly and quantitatively map FUS beams for quality assurance in the clinic, and to accelerate research and development of new FUS systems and techniques. However, conventional ultrasound pressure beam mapping instruments including hydrophones and optical techniques are slow, not portable, and expensive, and most cannot map beams at actual therapeutic pressure levels. Here, we report a rapid projection imaging method to quantitatively map FUS pressure beams based on continuous-wave background-oriented schlieren (CW-BOS) imaging. The method requires only a water tank, a background pattern and a camera, and uses a multi-layer deep neural network to reconstruct beam maps. Results at two FUS frequencies show that CW-BOS imaging can produce high-resolution quantitative projected FUS pressure maps in under ten seconds, that the technique is linear and robust to beam rotations and translations, and that it can accurately map aberrated beams.

## I. INTRODUCTION

Focused ultrasound (FUS) with pressures up to several megapascals (MPa) is a noninvasive therapeutic modality that has a broad range of established and emerging applications, including tumor and fibroid destruction, drug delivery, brain surgery^1^, blood-brain barrier opening and neuromodulation. Ablative FUS was recently FDA-approved for treating essential tremor^2^, and clinical trials are ongoing to establish its safety and efficacy in delivering Alzheimers disease drugs via blood brain barrier opening^3^. The method can be highly selective and can produce very sharp margins as narrow as six cells between an ablated lesion and viable tissue^4^. To maximize FUS’s therapeutic benefit, it is required to know how much acoustic energy is delivered and where it is delivered, with high spatial accuracy and precision. Furthermore, for therapeutic efficacy and safety it is necessary to assess whether the FUS system output changes between treatments, and to check for system failures which could dangerously alter energy delivery. Experts have recommended that rigorous quantitative beam mapping be performed on clinical systems two to three times monthly^5^. For these reasons, the ability to quantitatively map the acoustic beam in two or three spatial dimensions in the clinic is essential, and it is important for the safety and reproducibility of FUS treatments that instruments for rapid field characterization become available. FUS beam mapping is also essential for research and the development of new FUS technologies and techniques, such as new therapeutic transducers^6^, methods to propagate FUS beams through the skull and other bones^7–9^, acoustic lenses^10–12^, and FUS-transparent MRI RF coils^13^. Beam mapping is also essential for focused imaging transducers, whose mechanical index must be characterized to ensure they meet safety guidelines set by the US Food and Drug Administration.

The most widely-used FUS beam mapping instruments are hydrophones. They are ill-suited to rapidly mapping beams produced by FUS transducers, because they provide fine temporal resolution but (as illustrated in Fig. 1) they only sample one spatial location at a time, and a 3D motion stage must be used to move them through a tank to produce a spatially-resolved beam map. This results in long measurement times that even with variable density sampling schemes can take up to several hours for 3D volumes. Furthermore, fine temporal resolution is not required for the majority FUS applications where the transducer is operated in a continuous-wave mode. The long measurement times limit hydrophones’ usefulness in measuring beams at multiple power levels or across ranges of experimental variables. Common polyvinylidene fluoride (PVDF) hydrophones are not prohibitively expensive but can only measure sub-therapeutic pressure levels since they are easily damaged by cavitation. To overcome their speed and power limitations, hydrophone measurements have been combined with computational modeling (holography)^14,15^, but these methods still require a large number of hydrophone measurements over a two-dimensional surface. More expensive (> $10k USD) membrane^16^ and fiber optic hydrophones^17^ can withstand higher pressures, but they are less sensitive than PVDF hydrophones, and bandwidth limitations at high pressures can be a problem. Any instrument that sits in the focus will experience damage due to cavitation, and will require periodic repair and recalibration, and hydrophone systems lack the portability needed for clinical quality assurance measurements.

**FIG. 1.**
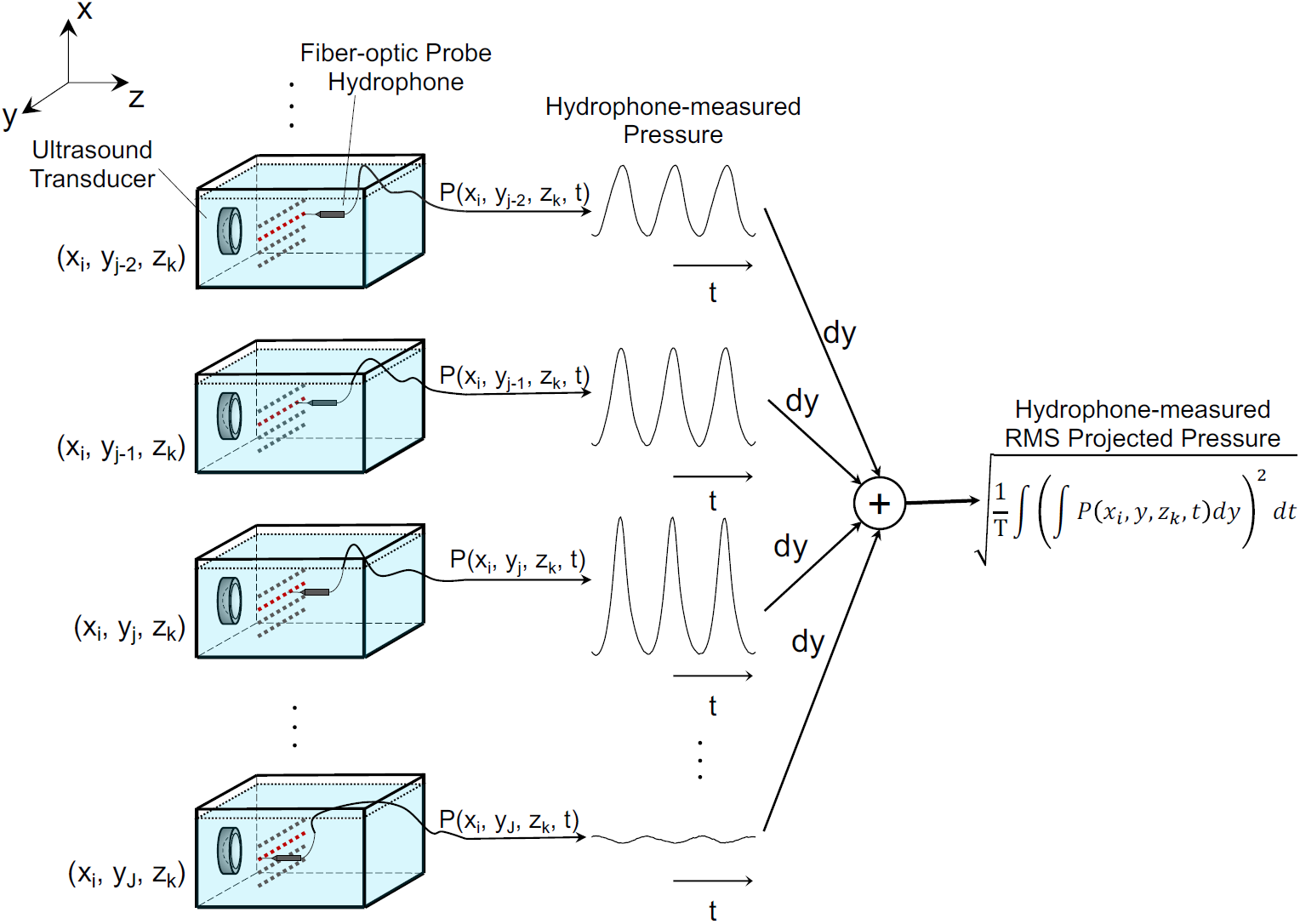
3D ultrasonic pressure field measurements using a hydrophone. The probe sensor of hydrophone samples only one spatial location at a time, so it must be translated by a motion stage to obtain a spatially-resolved map. To obtain the same root-mean-square (RMS) projected pressure map as the proposed CW-BOS method, hydrophone measurements are integrated along the line-of-sight dimension to obtain projected pressure waveforms, and then the RMS amplitude of the projected waveform is calculated.

Ultrasound pressure beams can also be mapped based on the deflection of light due to the acousto-optic effect. Optical ultrasound beam mapping methods such as photographic and laser schlieren methods have been used for more than fifty years to visualize ultrasound pressure fields in two dimensions^18–26^, and laser-based tomographic schlieren methods have been developed for temporally-resolved 3D ultrasound pressure field mapping^22,27–32^. The laser-based systems are based on the same physical principle as the technique proposed here, and are capable of impressive spatiotemporal resolution and sensitivity. However, they are limited to small FOVs and are prohibitively expensive (> $10k USD). Furthermore, to perform 3D mapping they typically require that the transducer itself be rotated, which makes them incompatible with in situ clinical transducers and limits their research utility.

Unlike conventional schlieren systems that require elaborate optical setups involving pulsed light sources, collimating lenses, and filters, background-oriented schlieren (BOS)^33^ imaging uses only a camera to image a background pattern through a nonuniform refractive index field. The background pattern is blurred by the nonuniform refractive index field, and cross-correlation of images acquired with and without the refractive index field in place produces index of refraction maps. In essence, in BOS the conventional sophisticated schlieren optical setup is traded for more sophisticated computation, which is much less expensive and easily replicated. The method has been used tomographically outside of acoustics to map static refractive index fields in 3D^33–39^, and it has been used to visualize FUS beams qualitatively in 2D^40^. However, the image formation process in BOS imaging of FUS beams is different from conventional BOS, because the refractive index is proportional to pressure^41^ which changes dynamically during a typical camera exposure time, so the background image is blurred rather than coherently displaced, and cross-correlation cannot be directly applied to extract refractive index or pressure maps. To freeze time to a fixed phase in the ultrasound cycle, tomographic BOS FUS beam mapping has been performed with a strobed light source^42,43^, but these methods have not yet been validated beyond qualitative comparisons to hydrophone measurements, and a strobed light source again complicates the setup and limits signal-to-noise.

Here we describe a rapid projection imaging method to quantitatively map FUS pressure fields based on continuous-wave BOS (CW-BOS) imaging. It requires only a water tank, a background pattern displayed on one side of the tank, and a camera to photograph the pattern through the other side of the tank (Fig. 2a). The proposed method leverages the recent availability of tablet PCs with high-resolution displays and consumer-grade digital single-lens reflex cameras with high pixel density that can resolve the sub-millimeter blurring of the BOS background pattern at a distance, as well as deep learning techniques that solve the difficult inverse problem of relating blurred photographs to projected pressure amplitudes. It can be implemented in a small and portable package to rapidly map FUS and focused imaging transducer beams in 2D, and there are no parts to experience wear from the FUS beam. Illustrated in Fig. 2b, the background images are bed-of-nails patterns, where each dot is blurred by the ultrasound beam in a distinctive pattern that can be interpreted as a histogram of local image displacement over time and is related to the projected root-mean-square (RMS) pressure field. Reconstruction is carried out using a deep neural network that relates each histogram in the photograph to an RMS projected pressure amplitude, and is trained from simulated photographs. Fig. 1 illustrates how the RMS projected pressure amplitude would be obtained using hydrophone measurements with a motion stage, which requires several hours of scan time, while the proposed method can produce the same measurement in seconds. The method was implemented and compared to fiber-optic hydrophone measurements, to evaluate its feasibility, accuracy and robustness.

**FIG. 2.**
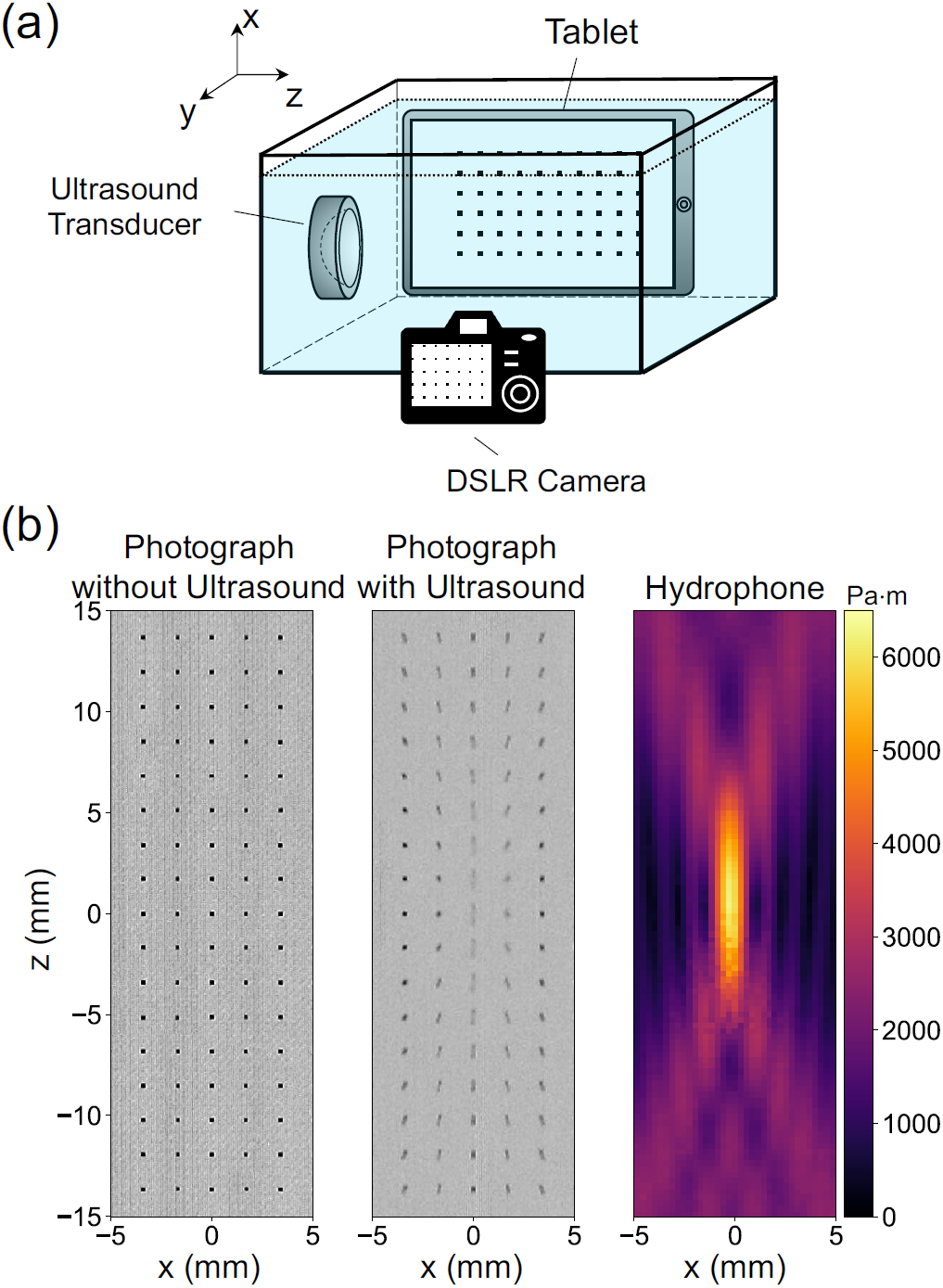
The proposed continuous-wave background-oriented schlieren (CW-BOS) beam mapping method displays a bed-of-nails pattern on one side of a water tank, and photographs it from the other side of the tank. When FUS is switched on, the nails blur in a distinctive pattern that can be related to the projected pressure field. a) A 2D CW-BOS system comprises a glass tank filled with water that is acoustically coupled to the ultrasound transducer, a tablet displaying a background pattern, and a camera to photograph the pattern when the FUS beam is turned on. b) Acquired photographs without and with FUS, and a hydrophone-measured RMS projected pressure map of the same FUS beam. In the photographs, the blurred nails are narrower and elongated along the beam propagation direction (top to bottom) in the focus, while the nails are blurred diagonally on either side of the focus. The FUS beam is propagating from top to bottom in these images.

## II. METHODS

### A. CW-BOS FUS Beam Mapping Hardware Setup

Fig. 3 shows the hardware setup for CW-BOS FUS beam mapping, which was built around an ultra-clear rimless water tank (Fragtastic Reef, Mankato, MN, USA) made of 5 mm-thick aquarium-grade glass. The size of glass tank was 31×19×19 cm^3^ (width × depth × height), and it was filled with degassed deionized water. To suppress reflections, an acoustic absorber (Aptflex F48, Precision Acoustic Ltd, UK) was placed against the tank wall opposite the FUS transducer. Two FUS transducers were used in this study: a 6.32 cm-diameter 1.16 MHz transducer with focal length 6.3 cm and f-number 2 (H101, Sonic Concepts, Bothell, WA, USA), and a 1.91 cm-diameter 2.25 MHz transducer with focal length 5.1 cm and f-number 2 (Valpey Fisher IL0206HP, Hopkinton, MA, USA).

**FIG. 3.**
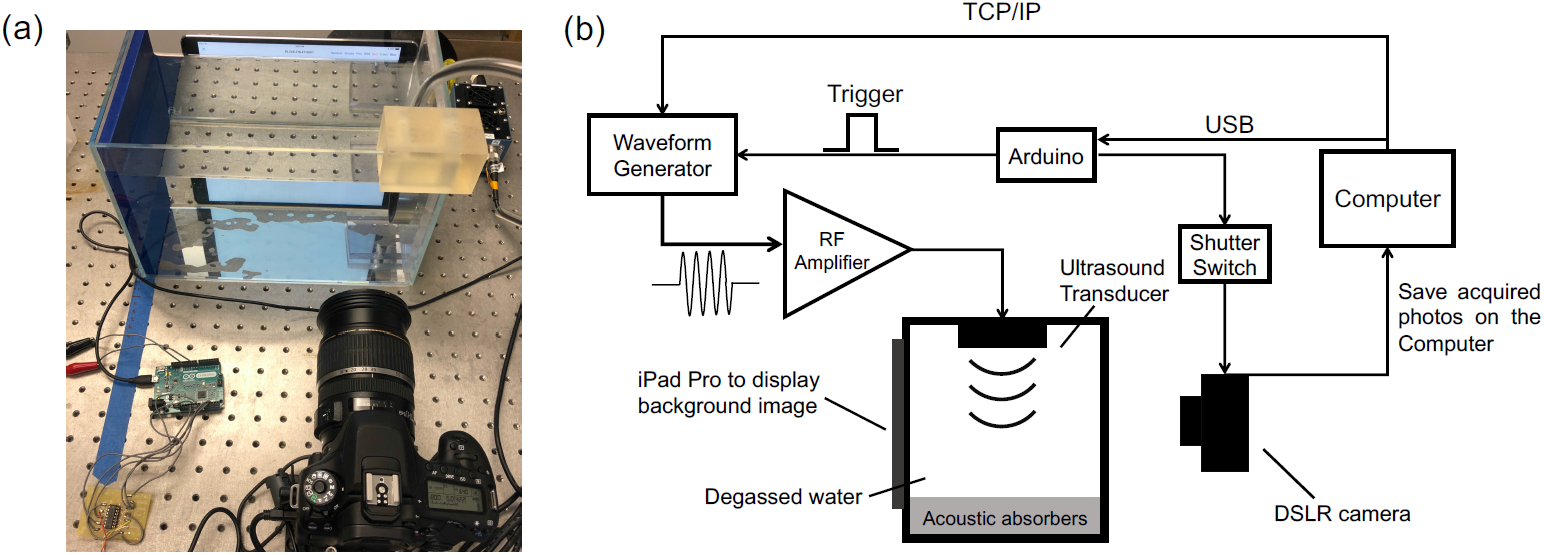
a) Photograph of the CW-BOS measurement setup used in this study. The camera is in the lower right and was centered on the nominal FUS beam focus. The iPad was placed against the opposite side of the water tank from the camera, the FUS transducer was mounted on the right side of the tank, and a blue acoustic absorber was mounted on the left side opposite the transducer to suppress reflections. b) Electrical diagram of the setup, including a top-down depiction of the tank. An Arduino was used to open the camera shutter a fixed time period after triggering the waveform generator, so that photos were taken when the FUS beam was at steady-state. The experiment was coordinated by a MATLAB script which set the waveform generator parameters and initiated an acquisition via the Arduino.

A 10.5” iPad Pro (Apple Inc, Cupertino, CA, USA) was placed against one of the long sides of the tank, which displayed bed-of-nails background images using a Python script running in the Pythonista app (OMZ Software, Berlin, Germany). The experiment computer told the iPad which background image to display via TCP/IP commands sent over WiFi. An EOS 80D 24.2 megapixel digital single-lens reflex (DSLR) camera with an EF-S 17-55mm f/2.8 IS USM lens (Canon Inc, Tokyo, Japan) was placed so that its body was 12 cm from the outer wall of the tank opposite the iPad, and was connected to the experiment computer via USB. The camera’s settings were controlled and photos were downloaded from it using the EOS Utility software (Canon Inc, Tokyo, Japan). An Arduino Leonardo R3 microcontroller board (Arduino, Italy) was used to open the camera shutter a fixed delay period after triggering the waveform generator, so that photos were taken when the FUS beam was at steady-state. The Arduino controlled the shutter via an analog switch (CD74HC4066E, Texas Instruments, Dallas, TX, USA) which electrically closed two switches (focus and shutter) of a modified wired manual shutter release that was connected to the N3 connector of the camera, and it sent a TTL trigger to the external trigger port of the FUS waveform generator (Keysight 33500B series, Santa Rosa, CA, USA) to initiate the FUS. The waveform generator’s parameters were set using TCP/IP commands sent from the computer via an ethernet connection, and its output was connected to an E&I A-150 amplifier (E&I Ltd, Rochester, NY, USA) to drive the transducer.

### B. CW-BOS Acquisition Details

Acquisitions were initiated by the experiment computer. When instructed by the computer, the Arduino sent a TTL pulse to the waveform generator to generate a 100,000-cycle, 86 ms pulse at 1.16 MHz, and a 150,000-cycle, 67 ms pulse at 2.25 MHz, then waited 50 ms and opened the camera shutter. The camera settings were: image size 4000×6000 pixels, ISO 640, shutter speed 1/800 s, f-number f/5. The photographs were saved on the computer in the RAW image format. The shutter speed corresponded to 1,450 FUS cycles for the 1.16 MHz transducer, and 2,813 FUS cycles for the 2.25 MHz transducer. During the experiments, the whole measurement setup was covered by a black cloth to suppress ambient light, so the iPad provided the only illumination.

The background images displayed by the iPad were bed-of-nails patterns comprising black dots/nails on a regular grid with a white background. The size of each dot was 2 pixels × 2 pixels. The distance between consecutive dots in each direction was 8 dots (16 pixels), which corresponded to a physical distance of 1.7 mm, and was set based on the maximum expected displacement in the experiments. To obtain a high-resolution beam map, 16 (4×4) photos were acquired across a series of equal-interval grid translations in the *x* and *z* dimension, as illustrated in Fig. 4. The images were segmented into small rectangular patches around each dot using MATLAB’s bwmorph function (Mathworks, Natick, MA, USA), then the RMS projected pressure was calculated by the neural network for each dot as described below, and those values were tiled into the final reconstructed beam map. With the camera placed a total distance of 31 cm from the iPad, each rectangular patch comprised between 42×42 and 46×46 pixels, and was upsampled to 54×54 pixels for reconstruction. To avoid optical color dispersion, only the green channel from the photos was used for reconstruction, which has the largest weight in Rec.ITU-R BT.601-7^44^. The total scan time for 5 averages was 3-5 minutes and was dominated by delays including photo transfers from the camera to the experiment PC. Without these delays, the total scan times were approximately 8 seconds (16 photos × 100 ms of FUS on-time × 5 averages).

**FIG. 4.**
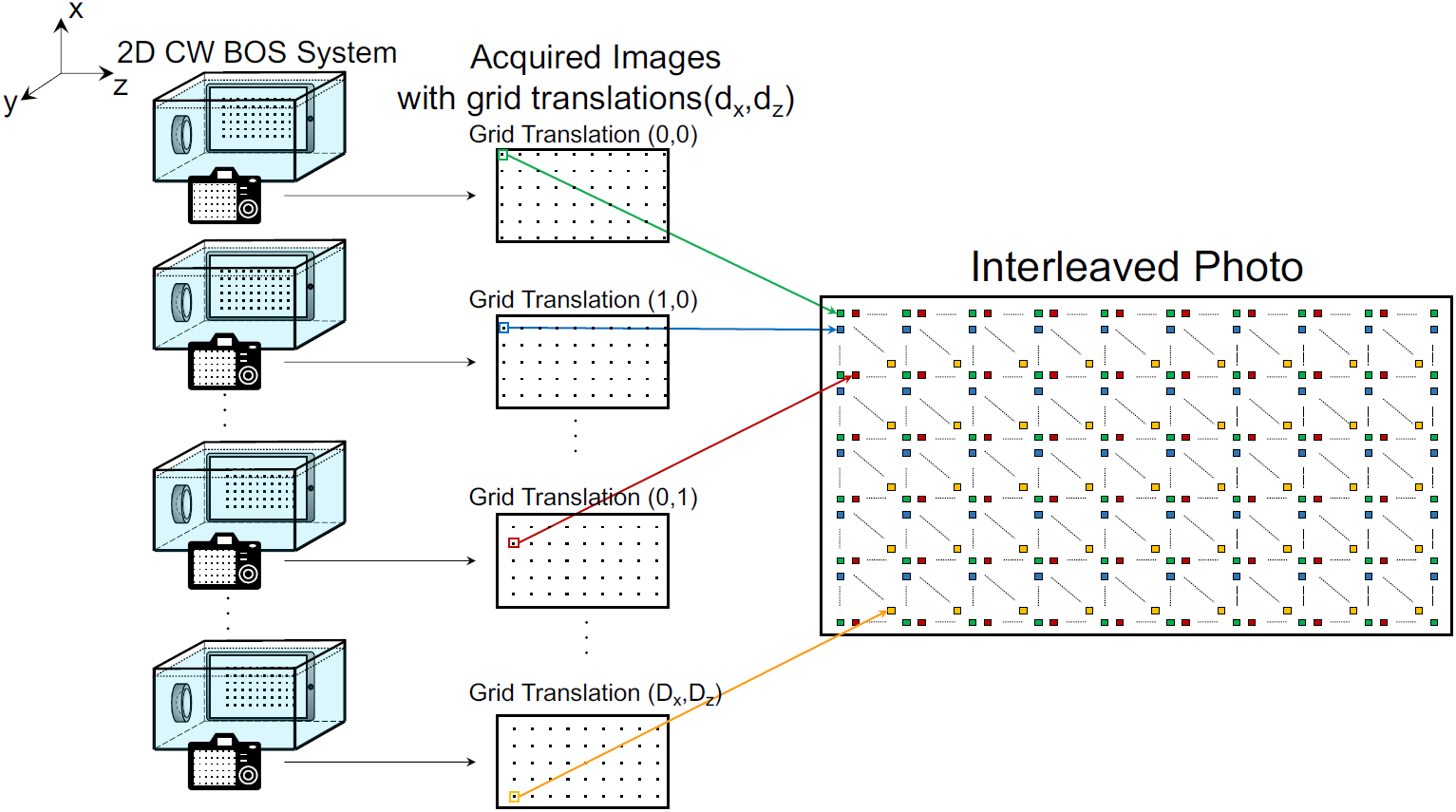
To measure a high-resolution beam maps, multiple CW-BOS images were collected over a range of grid translations in the *x* and *z* directions. The reconstructed projected RMS pressure values were then tiled into the final beam map.

### C. Optical Hydrophone Measurements

For validation, the FUS beams were also measured using fiber-optic hydrophones^45^ (136-10T and 132-03, Precision Acoustic Ltd, UK). The hydrophone measurements were performed in the same water tank as CW-BOS, using a Picoscope (Model 5242B, Pico Technology, UK) to record data to the computer, and a 3D motion stage (Image Guided Therapy, Bordeaux, France). The motion stage and Picoscope were both controlled by the experiment computer via USB. As illustrated in Fig. 1, to calculate reference RMS projected pressure maps from the hydrophone data, the synchronized hydrophone measurements were integrated along the line-of-sight (*y*) dimension to calculate the projected pressure waveforms and RMS projected pressure, using five FUS cycles from middle of the hydrophone-measured pulses. The 1.16 MHz maps were measured over a 10×10×30 (*x* × *y* × *z*) mm^3^ volume at 1.16 MHz, and a 10×10×47.5 mm^3^ volume at 2.25 MHz, with step sizes of 0.25 mm in *x* and *y*, and 0.25 mm in *z* at 1.16 MHz and 0.5 mm at 2.25 MHz. The total scan time was approximately 6.7 hours for the 1.16 MHz transducer (200,000 spatial locations) and 5.5 hours for the 2.25 MHz transducer (160,000 spatial locations).

### D. Mathematical Model for CW-BOS Imaging of FUS Pressure Fields

Fig. 2a shows that when a background image displayed by the tablet is photographed through the water tank by the camera, the image is distorted due to the refraction of light rays as they travel through the water from the tablet to the camera lens. The refraction angle in each of the photographed dimensions (*x* and *z*) is determined by the refractive index of the water:

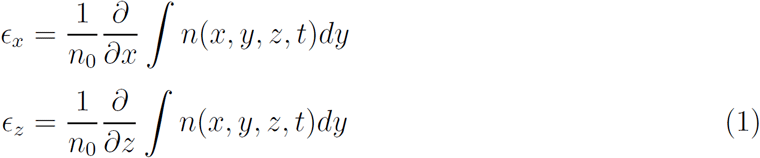

where *n*_0_ is the ambient refractive index of water, *n* is the 3D refractive index field, *y* is the projected (line-of-sight) dimension. The 3D refractive index field (*n*) is proportional to the acoustic pressure *p*^41^:

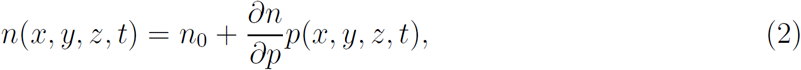

where 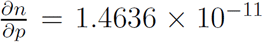 Pa^−1^ is the adiabatic piezo-optic coefficient^46,47^. Assuming the light ray is deflected as it passes through the FUS beam’s refractive index field and then continues across a distance *D* in the *y* dimension before being recorded by the camera, the image displacement at a location (*x, z*) in the photograph of the tablet’s image is obtained by substituting Equation 2 into Equation 1 and scaling by *D*:

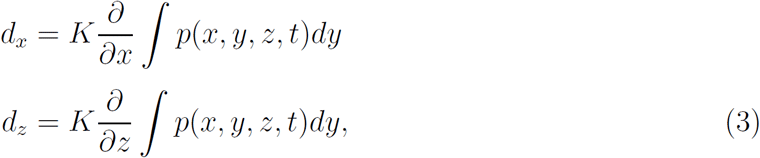

where 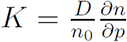. In this equation, the integral of the 3D pressure field along the projected dimension (*y*) is the projected pressure field *P*_*proj*_(*x, z, t*). This forward model was used to calculate the CW-BOS histograms to train the neural network reconstructor as described below.

### E. Numerical FUS Beam Simulations and Training Data Generation

To generate the FUS-blurred background images used to train the reconstructor, spatially- and temporally-resolved steady-state FUS pressure fields with nonlinearity were simulated using a modified angular spectrum method^48^ with frequency domain attenuation and dispersion, absorbing boundary layers, and an adaptive propagation step size. An operator splitting term was used to separate the terms in the retarded-time formulation of the nonlinear angular spectrum equation^49^. The attenuation and dispersion term was solved directly in the frequency domain using a filtering approach. The nonlinear term was solved in the time domain using a Rusanov scheme to accurately capture the shock front in a flux-conservative fashion^50^. The simulations used a speed of sound of *c*_0_ = 1500 m/s, *λ/*8 grid spacing in the dimensions transverse to beam propagation (0.16 mm at 1.16 MHz and 0.08 mm at 2.25 MHz), *λ/*4 grid spacing in the axial/propagation dimension (0.32 mm at 1.16 MHz and 0.16 mm at 2.25 MHz), a nonlinearity coefficient of *β* = 3.5, an equilibrium density *ρ*_0_ = 1000 kg/m^3^, and a dwell time of 1/(40*f*_0_) (21.5 ns at 1.16 MHz and 11.1 ns at 2.25MHz). The beams were simulated over a 9.5×9.5×9.5 cm^3^ volume for the larger 1.16 MHz transducer and a 3.8×3.8×3.8 cm^3^ volume for the smaller 2.25 MHz transducer. The simulated transducers generated 12-cycle pulses, and the middle cycle was saved at each spatial location, representing steady-state. A total of 34 simulations were run for the 1.16 MHz transducer, for peak negative pressure amplitudes between 1 - 11.5 MPa, and f-numbers of 1 and 2. A total of 38 simulations were run for the 2.25 MHz transducer, for 1.2-10 MPa, and f-numbers of 1 and 2.

The training data for the reconstructor comprised histograms paired with their projected RMS pressure values. First, projected pressure waveforms were calculated by integrating the beams along the *y* dimension, and projected RMS pressure values were calculated from those waveforms. To calculate a histogram for each simulated (*x, z*) location, projected pressure waveforms were first calculated, then image displacements were calculated using Equation 3, which required finite differencing the *y*-projected pressure fields in the *x* and *z* dimensions and scaling the result by the distance *D* between the focus and the tablet screen (*D* = 8.5 cm for our hardware setup). Then, for each time instant, a distorted image was computed by shifting the spatial location’s nail by the calculated image displacements in each direction, and the distorted images were summed over one ultrasound period to obtain a final simulated BOS histogram image. The simulated histograms were convolved with a point spread function measured from an undistorted (no-FUS) photograph taken with our system. The histograms were individually normalized for zero mean and unit standard deviation, and the RMS projected pressures were collectively normalized. The histogram dimensions were 54×54, which corresponded to a spatial area of size 1.7 × 1.7 mm^2^ on the screen of the tablet, with a pixel width of 0.024 mm. The 1.16 MHz training data comprised a total of *N* = 744885 examples, and the 2.25 MHz training data comprised a total of *N* = 554905 examples.

Prior to inputting them to the reconstructor network, the training histograms were compressed to a dimensionality smaller than their number of pixels by projecting them to a subspace derived by singular value decomposition (SVD) truncation^51,52^. For each FUS frequency, a dictionary was formed from all the training data by reshaping the histograms to length-*M* row vectors **d** ∈ *R*^1*×M*^, where *M* = 54^2^ = 2916, and stacking them into a dictionary matrix **D** ∈ *R*^*N×M*^, where *N* is the number of training examples. The matrix **D** was decomposed by SVD into the product of three matrices, **D** = **USV**^*T*^, where **U** ∈ *R*^*N×M*^ is an orthonormal matrix containing the left singular vectors, **S** ∈ *R*^*M×M*^ is a diagonal matrix containing the singular values, and **V** ∈ *R*^*M×M*^ is an orthonormal matrix of right singular vectors. A lower dimensional compressed subspace was obtained by truncating the SVD to its first *K* singular values (where *K* = 141 at 1.16 MHz and 118 at 2.25 MHz), and the vectors of the resulting truncated right singular vector matrix **V**_*K*_ ∈ *R*^*M×K*^ spanned this lower-dimensional subspace. Thereafter, each training and experimental BOS histogram **d** was projected to the lower-dimensional subspace by multiplying it with the matrix **V**_*K*_ to obtain its compressed coefficients **c** = **dV**_*K*_. These coefficients were the inputs to the neural network to obtain the projected RMS pressure values, as illustrated in Fig. 5a. The matrices **V**_*K*_ were stored and used to compress experimentally measured histograms prior to reconstruction of their projected RMS pressures.

**FIG. 5.**
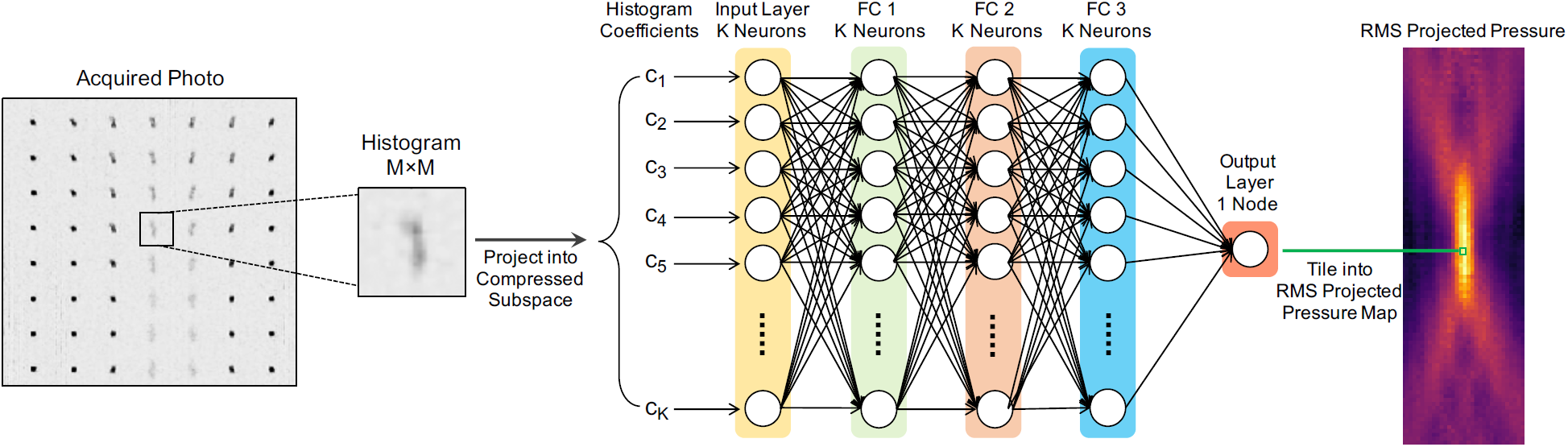
Schematic representation of neural network-based pressure reconstruction for each photographed histogram. Each histogram in a photo is segmented into an *M* × *M* sub-image, and then projected to the compressed subspace, and its *K* (*K* ≪ *M*) coefficients in that subspace are input to a deep neural network with a fully-connected input layer and three fully-connected hidden layers and hyperbolic tangent activations, which feed a single-node layer that outputs the final estimated RMS projected pressure value, which is tiled into the final beam map.

### F. Neural Network Architecture and Training

To reconstruct RMS projected pressure maps from a set of photographs, Fig. 5 shows that each histogram is projected into the compressed SVD subspace, and the resulting length-*K* vector of coefficients **c** is input to a deep neural network comprising three fully connected layers (FC1 to FC3 in Fig. 5). The input and fully connected layers all have *K* nodes, and each is followed by a hyperbolic tangent activation function. The output layer comprises a linear activation function and has one node, the output of which is the RMS projected pressure for the input histogram coefficients.

A network was trained for each frequency in Keras^53^ on the Tensorflow deep learning framework^54^ using two NVIDIA graphics processing units (NVIDIA, Santa Clara, CA, USA) for 20 epochs (approximately 1 hours), on Vanderbilt University’s parallel computing cluster (Advanced Com-puting Center for Research and Education, Vanderbilt University, Nashville, TN). Each batch was trained for 400 steps. The optimization algorithm RMSProp^55^ was used with mini-batches of size 1024, a learning rate of 0.00005, momentum 0.0 and decay 0.9. Mean squared error was used as the loss function for training. An additional *L*_1_-norm penalty (*λ*_1_ = 0.00002) was used to promote sparsity of the weights in the input layer, and an *L*_2_-norm penalty (*λ*_2_ = 0.0002) was used to prevent overfitting in the each layer. Keras’s real-time augmentation was used to rotate the training histograms by 0°-30° to achieve robustness to transducer rotations. Given all the acquired photos, the final beam map reconstructions took approximately 20 seconds of computation on a desktop computer with a 4.2 GHz Intel Core i7 CPU and 32 GB 2400 MHz DDR4 RAM (iMac, Apple Inc, Cupertino, CA, USA).

## III. RESULTS

### A. Two FUS Frequencies

FUS frequencies range from hundreds of kHz to several MHz. To demonstrate the CW-BOS method at different frequencies, mapping was performed for 1.16 and 2.25 MHz transducers and compared to optical hydrophone measurements. The FUS pulses were produced by waveform generators with voltage amplitudes of 200 millivolts peak-to-peak (mV_pp_) (1.16 MHz) and 100 mV_pp_ (2.25 MHz), which corresponded to peak negative pressures (PNP) of −4.5 MPa (1.16 MHz) and −1.4 MPa (2.25 MHz), as measured by the optical hydrophone. Fig. 6a shows the hydrophone-measured RMS projected pressure map (left) and reconstructed CW-BOS RMS projected pressure map (right) at 1.16 MHz, where CW-BOS reconstruction was performed by segmenting the blurred photograph into a patch containing each nail and then inputting each segmented patch to the deep neural network to obtain the RMS projected pressure at that point. The amplitudes and shapes of the hydrophone and CW-BOS beams matched closely, with a root-mean-squared error (RMSE) of 298 Pa·m, or 4.8% of the hydrophone-measured peak amplitude, and main-lobe full-width at half-maximums (FWHM’s) of 1.5 mm (hydrophone) versus 1.4 mm (CW-BOS) in the *x* dimension, and 11.4 mm (hydrophone) versus 11.7 mm (CW-BOS) in the *z* dimension. Fig. 6b shows the hydrophone (left) and CW-BOS (right) RMS projected pressure maps at 2.25 MHz. The RMSE between the two was 192 Pa·m, or 7.9% of the hydrophone-measured peak amplitude, the main-lobe FWHM’s in *x* were 2.1 mm (hydrophone) versus 2.0 mm (CW-BOS), and the main-lobe full width at 80% of maximum’s in *z* were 26.4 mm (hydrophone) versus 25.9 mm (CW-BOS).

**FIG. 6.**
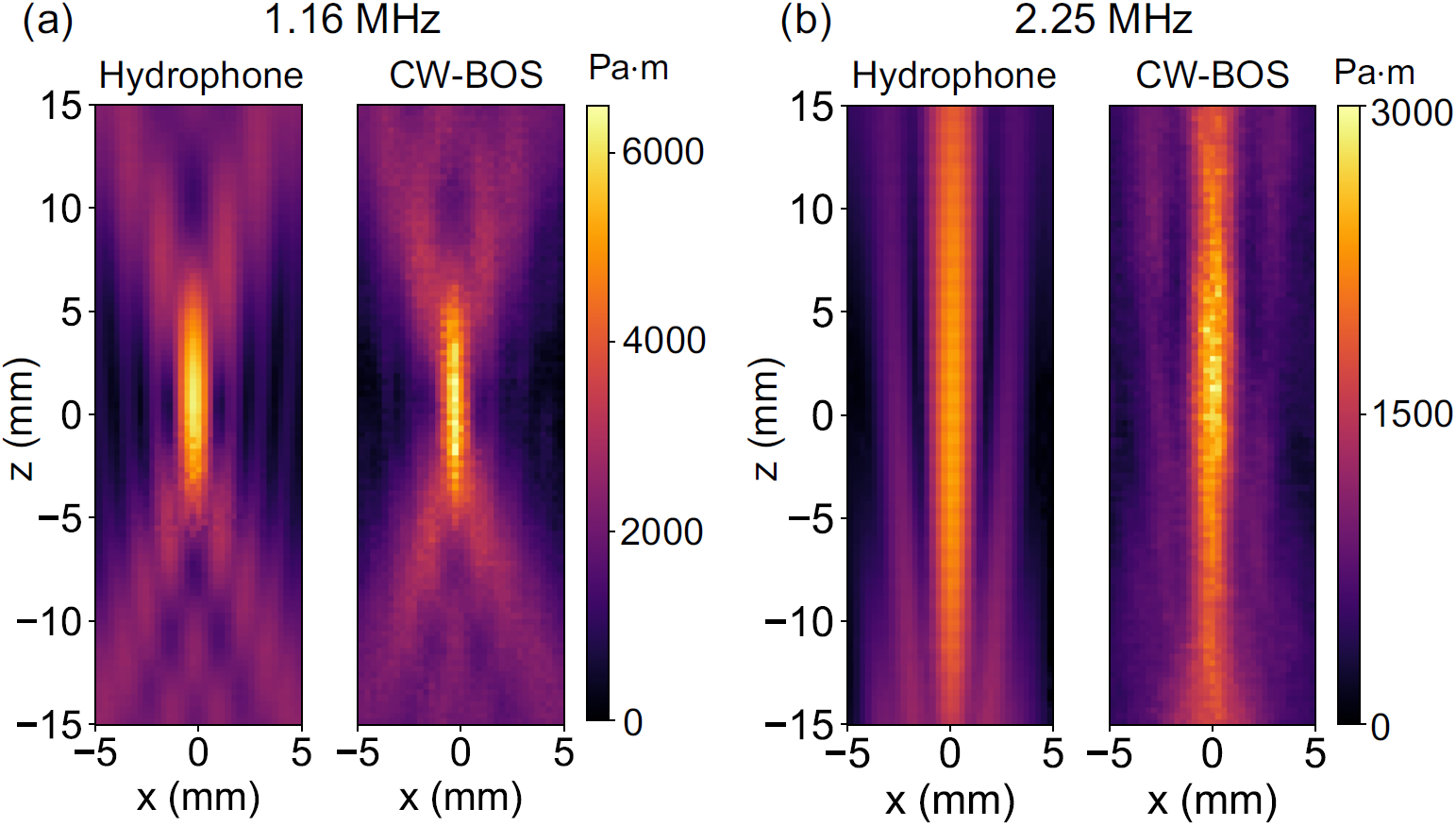
Comparison of projected pressure maps measured using an optical hydrophone and CW-BOS for two transducers at different frequencies. a) Maps measured for a 1.16 MHz transducer driven at 200 mV_pp_. b) Maps measured for a 2.25 MHz transducer driven at 100 mV_pp_.

### B. Linearity

CW-BOS images were further acquired across transducer driving voltage amplitudes. Data were acquired with the 1.16 MHz transducer and waveform generator amplitudes between 0 and 200 mV_pp_, in 25 mV_pp_ steps. Fig. 7a shows reconstructed projected pressure maps across driving voltage amplitudes. Fig. 7b plots the mean across five repetitions of the projected pressure in the focus (indicated by the arrow in Fig. 7a) at each amplitude, along with optical hydrophone measurements which were taken at 50, 100, 150, and 200 mV_pp_, which corresponded to PNPs of −1.5 MPa (50 mV_pp_), −2.5 MPa (100 mV_pp_), −3.6 MPa (150 mV_pp_), and −4.5 MPa (200 mV_pp_). The error bars represent the standard deviation of the values over the five repetitions. The fitted slopes of the RMS projected pressure amplitudes were 32.2 Pa·m/mV_pp_ (hydrophone) versus 32.3 Pa·m/mV_pp_ (CW-BOS). The Pearson’s r-value between the hydrophone and CW-BOS measurements was 0.998.

**FIG. 7.**
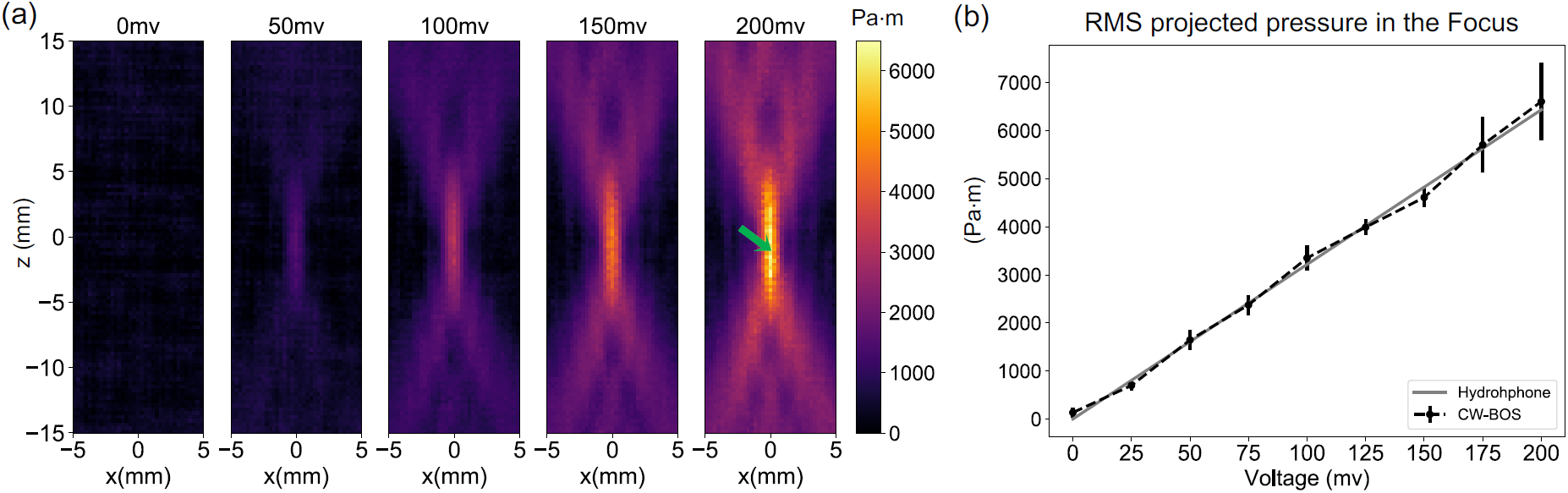
CW-BOS pressure maps versus driving voltage amplitude. a) Reconstructed RMS projected pressure maps for the 1.16 MHz transducer. b) RMS projected pressure at the focus measured by the hydrophone and CW-BOS, where the CW-BOS values were averaged across five repeated measurements, and the error bars represent standard deviation across the measurements.

### C. Signal-to-Noise Ratio and Number of Averages

Fig. 8 shows that noise can be reduced by averaging reconstructions from repeated CW-BOS acquisitions. Fig. 8a shows reconstructed CW-BOS projected pressure maps between one and eight averages for a driving voltage of 200 mV_pp_, and the apparent noise is reduced significantly comparing 1 and 8 averages. Ten repetitions were further acquired at driving voltages of 50, 100, 150, and 200 mV_pp_, and Fig. 8b plots the decremental RMS error in a 5.6×20 mm^2^ region centered on the focus between maps reconstructed from one to ten averages. The differences stopped changing significantly after five averages. Fig. 8c plots the incremental signal-to-noise ratio (SNR) around the focus of reconstructed 200 mV_pp_ maps from one to ten averages; with one acquisition the SNR was 40, but was improved to 70 by five averages. Here, SNR was calculated as the ratio of the signal amplitude in the middle of the focus to the standard deviation in background regions without significant projected pressures. Overall, for our setup, the maps stop changing significantly after five averages, so this number was used for all experimental results.

**FIG. 8.**
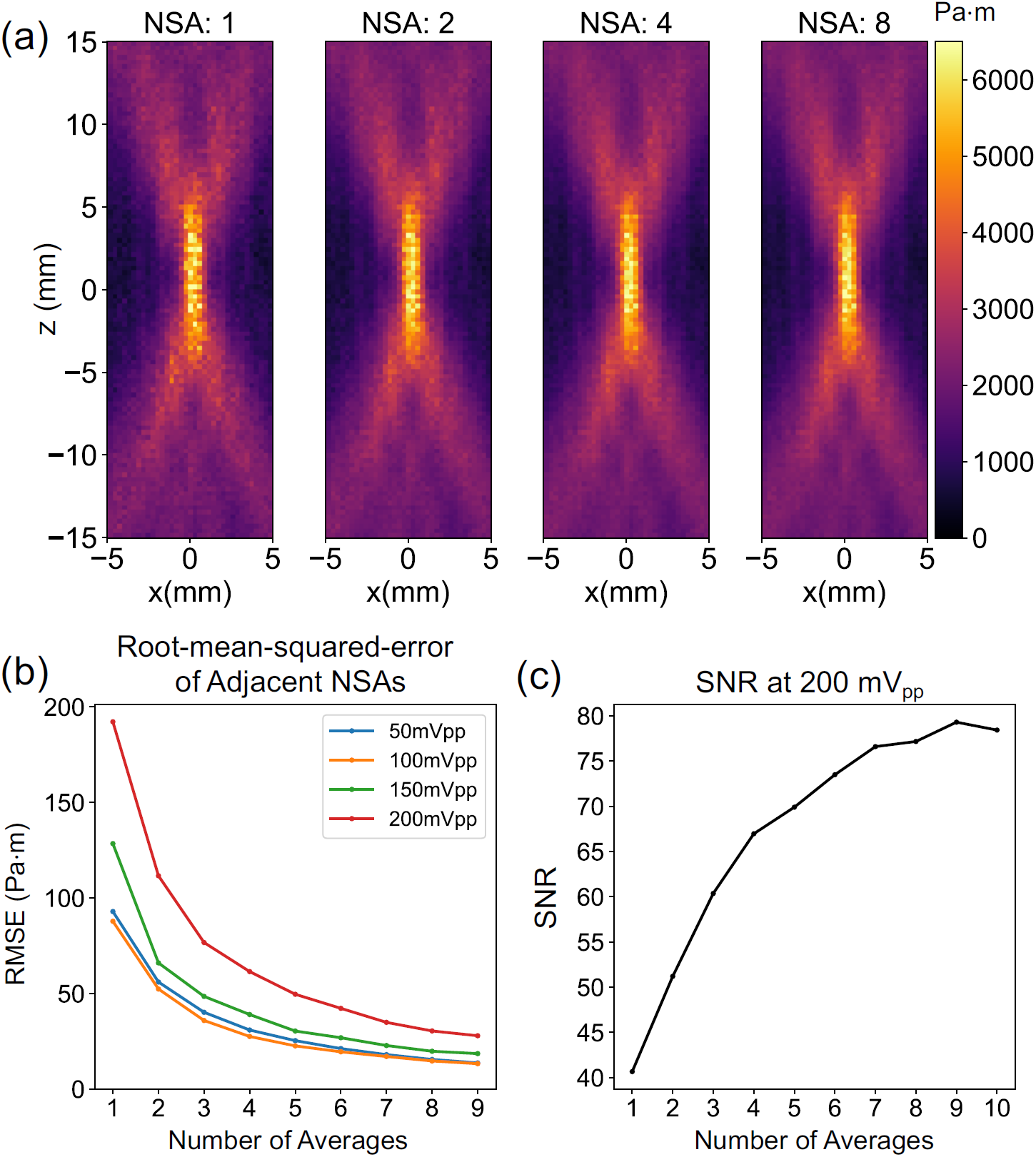
Signal-to-noise and reconstruction from different numbers of averages. a) Reconstructed RMS projected pressure maps with 1 to 8 averages with a driving voltage of 200 mV_pp_ at 1.16 MHz. b) Mean-squared-error between maps resulting from *i* + 1 versus *i* averages in a 5.6 × 20 mm^2^ region around the focus. c) SNR around the focus with a driving voltage of 200 mV_pp_ and one to ten averages.

### D. Rotational and Translational Invariance

User error could introduce rotations and displacements between the camera and the FUS beam, so the method should be robust to a reasonable range of such errors. The top row of Fig. 9a shows CW-BOS RMS projected pressure maps imaged with a 200 mV_pp_ driving voltage, with no transducer rotation (the reference case), and 15°and 30° rotation about the y (line-of-sight) axis. The shape and intensity of the rotated pressure fields are unchanged compared to the reference, because the reconstruction was trained with rotated beam maps to accommodate rotations. The projected pressure amplitudes in the focus were 5934 Pa·m (0°), 6152 Pa·m (15°) and 5749 Pa·m (30°). FWHM’s in the *x* dimension were 1.4 mm (0°), 1.7 mm (15°) and 1.8 mm (30°), and 11.7 mm (0°), 10.8 mm (15°), and 11.3 mm (30°) in the *z* dimension. Fig. 9b further shows CW-BOS RMS projected pressure maps measured with the camera translated ±2.5 cm along the z-dimension, with a 150 mV_pp_ driving voltage. The intensity and shape of pressure fields were again unchanged compared to the reference. The projected pressure amplitudes around the focus were 4348 Pa·m (no translation), 4616 Pa·m (−2.5 cm) and 4344 Pa·m (+2.5 cm). FWHM’s in the *x* dimension were 1.5 mm (no translation, −2.5 cm and +2.5 cm), and 12.1 mm (no translation), 11.5 mm (−2.5 cm), and 11.4 mm (+2.5 cm) in the *z* dimension.

**FIG. 9.**
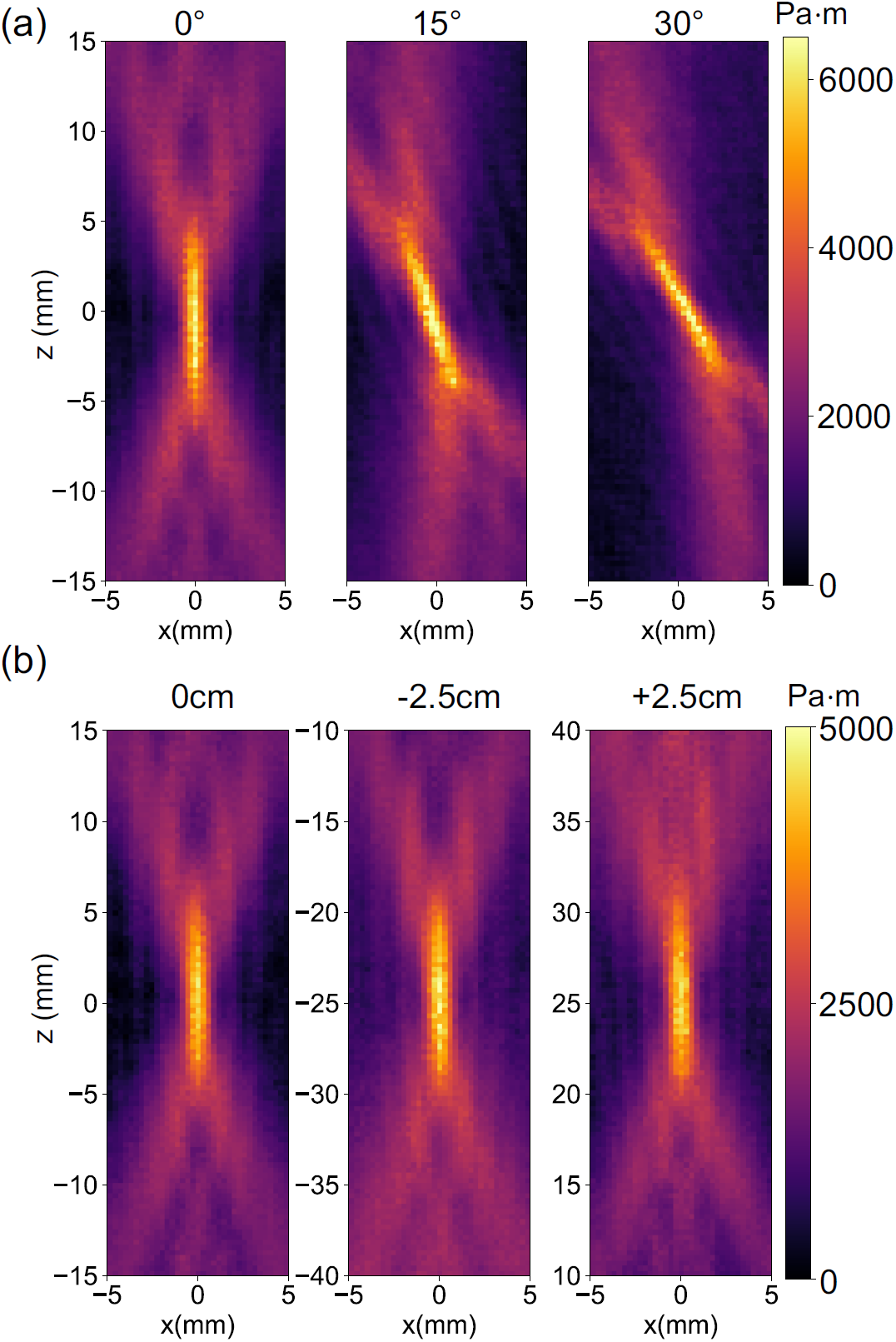
Rotational and translational invariance. a) Reconstructed RMS projected pressure maps obtained by rotating the 1.16 MHz transducer 0°, 15°, and 30°. b) Reconstructed RMS projected pressure map obtained with the camera focus centered on the focus, and shifted ±2.5 cm along the z-axis.

### E. Aberrations

An important potential application of the CW-BOS projection beam mapping method is to detect beam aberrations on clinical FUS systems. Fig. 10a shows an acoustic aberrator^56^ made from silicone (Elite double 8, Zhermack, Badina Polesine, Italy) constructed to block the bottom half of the 1.16 MHz transducer. CW-BOS RMS projected beam maps and hydrophone beam maps measured with this aberrator configuration are shown in Fig. 10b. The beams’ intensities and shapes are closely matched, and the RMSE between them was 256 Pa·m (10.8% of the hydrophone-measured peak amplitude). Fig. 10c further shows the lens placed to block the left half of the transducer, and Fig. 10d shows measured beam maps with this configuration. The maps again correspond closely, with an RMSE of 442 Pa·m (22.6% of the hydrophone-measured peak amplitude).

**FIG. 10.**
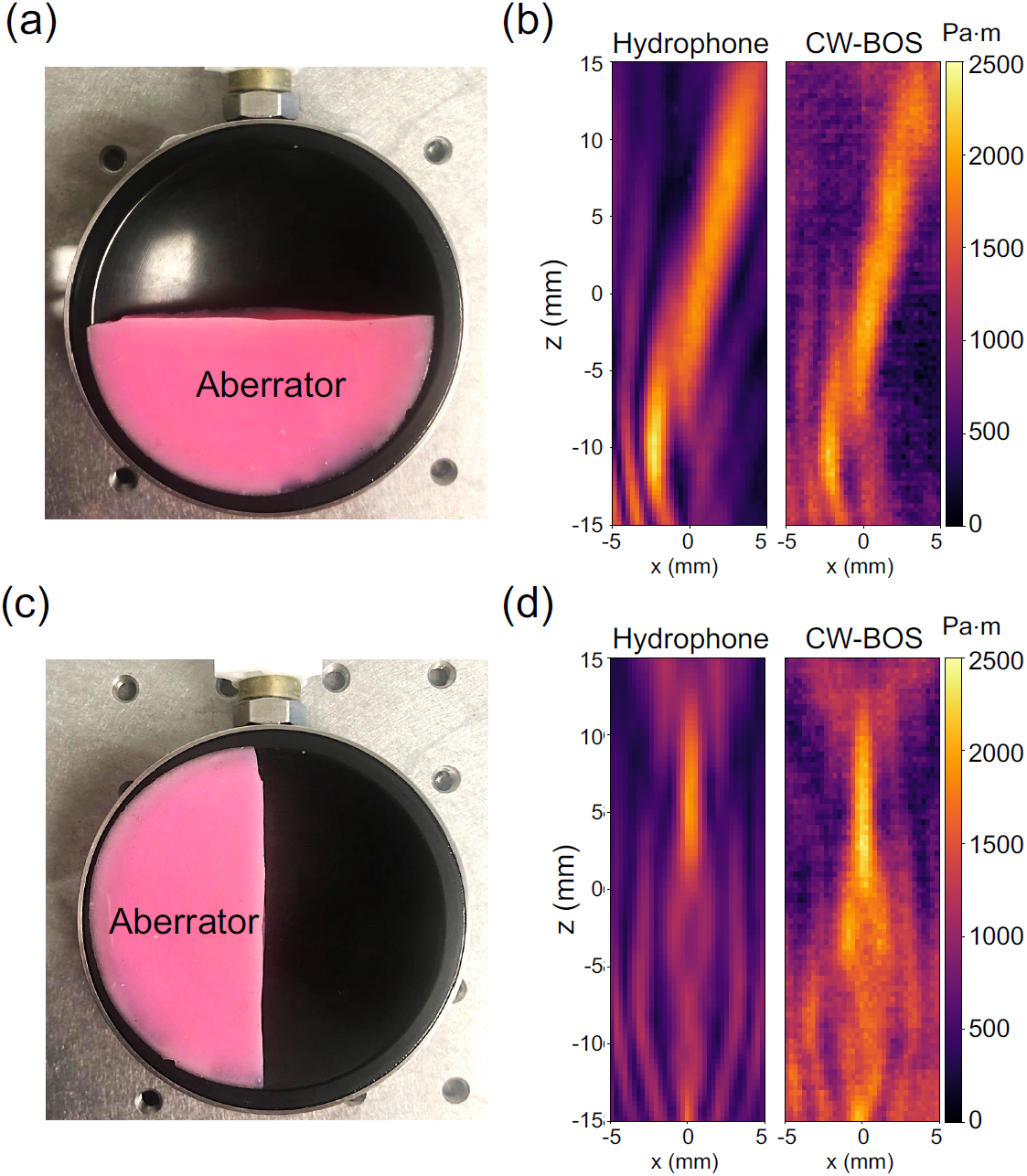
Aberrated beam mapping. a) An aberrator made from silicone was placed in front of the bottom half of the 1.16 MHz transducer. b) Optical hydrophone and CW-BOS projected beam maps measured with the bottom half of the transducer blocked. c) The aberrator positioned to block the left half of the transducer. d) Optical hydrophone and CW-BOS projected beam maps measured with the left half of the transducer blocked.

## IV. CONCLUDING REMARKS

Quantitative and fast mapping of FUS pressure fields is essential for treatment planning, safety, dosimetry, quality assurance and technical research^5^. We have proposed and demonstrated a rapid and inexpensive optical method to quantitatively map FUS pressure fields in two dimensions, which requires only a water tank, a tablet to display background patterns, a camera and a PC to reconstruct beam maps. The method could also be used to map the beams of focused imaging transducers, which are capable of generating similar pressure amplitudes to the FUS transducers evaluated here; the described experimental setup was sensitive to pressure beams with peak negative pressures less than 1 MPa. Unlike previous optical beam mapping methods that used strobed light sources to “freeze” the ultrasound beam at different phases so that it can be reconstructed algebraically^42^, we simplified the hardware setup by allowing the beam to run continuously during the acquisition which causes a blur rather than a coherent shift of the background pattern, and then used a deep neural network to solve the difficult inverse problem of reconstructing projected pressure amplitudes from the blurred image at each location in the photograph. We described a complete 2D BOS hardware system and acquisition protocol, described a forward model for image formation, and established a reconstruction. It is important to note that the reconstruction network operates only on one spatial location at a time, and does not make assumptions about spatial smoothness or structure of the beam in the imaged 2D plane, yet it produced beam maps that closely matched optical hydrophone measurements. This way, the technique maintains generality for important applications where beam structure would be difficult to predict, such as when mapping aberrated fields as was demonstrated here, or when the beam rotates or moves.

CW-BOS is an inexpensive (under $2000 USD) and rapid 2D FUS beam mapping tool based on a consumer-grade tablet and camera, with no moving parts or parts that can experience wear from the FUS beam. To make it portable, the tank could be sealed, the tablet and camera could be rigidly attached to it, and FUS could be coupled into it via a mylar membrane. Sealing the tank could also enable replacement of degassed water with a transparent liquid or gel that better approximates ultrasound propagation and absorption in tissue. This would represent the first truly portable beam mapping method. The reconstructor network would need to be trained for the specific frequency of the transducer, though with further work it may be possible to train a single network for a wide range of FUS frequencies. Our total CW-BOS scan times were 3-5 minutes, which was dominated by delays including photo transfers from the camera to the PC, and comprised less than 8 seconds of FUS-on time. We expect that with optimization, the total scan duration could be reduced to less than 10 seconds; reconstruction in MATLAB and Python then took another 20 seconds which could be further optimized, and does not require a high-end computer. Overall, the method achieves an approximate 2000x speedup compared to the time required to obtain the same information using a hydrophone. While the proposed hardware is not currently compatible with very large-aperture transcranial FUS transducers whose foci do not extend beyond their shell, it may be possible to map these systems by projecting background patterns onto the transducer surface.

There are several possible ways to improve or extend the proposed technique. First, a larger convolutional neural network that operates on entire photos rather than individual segmented histograms may achieve improved accuracy by learning spatial relationships between blurring patterns and FUS beam features, and it could enable the use of a single, dense background pattern to reduce acquisition times. However, this would require a much larger training corpus to maintain generality, as well as more computation and memory, both for training and reconstruction. We also assumed a parallel ray geometry between the background pattern and the camera in this work, but it may be possible to generate more accurate training histograms using ray tracing^57,58^. Finally, it may be possible to extend the method to reconstruct not just projected waveform amplitudes but also projected waveforms themselves, which would be needed to reconstruct 3D beam maps. A 3D system would also require the optics and the transducer to be rotated with respect to each other.

## ACKNOWLEDGMENTS

This work was supported by NIH grant R21 EB 024199. The authors would like to thank Charlotte Sappo for help with the shutter switch, and Marshall (Tony) Phipps with help using the optical hydrophone and FUS amplifier.

